# GNN-SubNet: disease subnetwork detection with explainable Graph Neural Networks

**DOI:** 10.1101/2022.01.12.475995

**Authors:** Bastian Pfeifer, Afan Secic, Anna Saranti, Andreas Holzinger

## Abstract

The tremendous success of graphical neural networks (GNNs) has already had a major impact on systems biology research. For example, GNNs are currently used for drug target recognition in protein-drug interaction networks as well as cancer gene discovery and more. Important aspects whose practical relevance is often underestimated are comprehensibility, interpretability, and explainability. In this work, we present a graph-based deep learning framework for disease subnetwork detection via explainable GNNs. In our framework, each patient is represented by the topology of a protein-protein network (PPI), and the nodes are enriched by molecular multimodal data, such as gene expression and DNA methylation. Therefore, our novel modification of the GNNexplainer for model-wide explanations can detect potential disease subnetworks, which is of high practical relevance. The proposed methods are implemented in the GNN-SubNet Python program, which we have made freely available on our GitHub for the international research community (https://github.com/pievos101/GNN-SubNet).

## I. Introduction

Graph Neural Networks (GNNs) have attracted much attention in general [1], [2], and in bioinformatics [3] and biomedical research in particular [4].

Recently, significant research efforts have been made to apply Deep Learning (DL) methods to graphs [5]. This progress resulted in useful advances in graph analysis techniques, which are useful in many biomedical applications [6], e.g. to use graph-based deep learning models for protein-drug interaction detection [7].

A very recent work presents graph-based frameworks for detecting novel cancer genes by using GNNs for node classification [8]. The authors label genes of a protein-protein interaction (PPI) network according to their cancer relevance (relevant or not). DL-based node classification is applied to predict whether or not unlabeled proteins are relevant to cancer.

A key feature of GNNs is that they enable the integration of knowledge graphs [9] into the algorithmic pipeline, such as ontologies [10], [11] and/or PPI networks [12], [13]. This feature allows a domain expert to integrate human experience, human conceptual knowledge and contextual understanding into the machine learning architectures. Such a human-in-the-loop (expert-in-the-loop) [14] can sometimes - of course not always - help to obtain more robust, reliable and also more interpretable results [15] [16]. It should be emphasized that robust, explainable, and trustworthy solutions are among the major goals of the AI community for the near future [17].

Such solutions are of practical relevance in critical areas where we suffer from low data quality, especially where we just do not have the i.i.d. data we actually need. Therefore, the use of AI in areas that impact human life (e.g. agriculture, climate, health, …) has led to an increased demand for trustworthy AI. This is especially true in sensitive areas such as biomedicine, where traceability, transparency and interpretability are not ends in themselves, but are now even mandatory due to regulatory requirements [18]. Finally, the “why” [19] is often more important to science than a pure result. Consequently, both explainability and robustness can promote reliability and trust and ensure that humans remain in control and thus that human intelligence is supported by artificial intelligence and by no means replaced [20].

In our work, we place a particular emphasis on the integration of PPI networks for disease subnetwork detection. Most existing methods for disease subnetwork detection rely on unsupervised clustering and/or community detection algorithms to detect modules with correlated node features.

We believe that functional subnetworks [21] with high classification accuracy, where node features are not necessarily correlated, may contain an additional set of biologically relevant disease modules. While conventional feature selection methods can be used for this task, most of them are not directly applicable to graph-structured data. This is where we come in, as an exception is our proposed method [22], where we introduce a greedy decision forest for subnetwork detection. To demonstrate the applicability of this approach, we enriched the nodes of a PPI network with multi-omic features. Decision trees are derived from this network using random walks. The decision trees evolve on this network to a minimal set of high-performance subnetworks. In this work, while we pursue a similar research goal, we further use powerful graph deep learning architectures and explainable AI methods [23] for DL-based disease subnetwork detection. To the best of our knowledge, this is novel and thus represents the first work that uses explainable AI for disease subnetwork discovery.

This paper is organized as follows: First, a brief summary of the proposed methodology for disease subnetwork detection is given. Section 3 describes the methodology, the GNN methods used and the validation data in detail. Section 4 presents the results obtained using synthetic datasets as well as multimodal human cancer data. Section 5 discusses the work presented and possible future research directions.

## II. New Approach

In this work, we propose explainable GNNs for the detection of disease subnetworks. We have formulated the subnetwork detection task as a graph classification problem, where graph topologies are the same for all instances, but the node feature values vary. The following methodology is presented.

Each patient is represented by the graph topology of a PPI network, where proteins are reflected by the nodes and the edges indicate a functional relationship between these proteins. The nodes of the patient-specific graphs are enriched by multiomic feature values, such as mRNA gene expression and DNA methylation. Following, we perform graph classification in order to classify patients into a cancer-specific group and a randomized cancer group. As a consequence, a GNN model is trained on domain-knowledge induced trajectories, which may result in more reliable and interpretable outcomes [24].

In order to ultimately uncover the decisions of the GNN classifier, we utilize a modified version of the GNNexplainer algorithm, by optimizing a *model-wide* node feature mask (see ‘Materials and Methods’ section for details). From the obtained node importance values we compute edge relevant scores. We assign these values as edge-weights to the PPI Network and apply weighted community detection algorithms. The detected communities with high edge importance scores represent the potential disease subnetworks.

## III. Materials and Methods

### A. GNN architecture

We have employed a GNN classifier as evaluated and implemented by [25]. The authors propose a Graph Isomorphism Network (GIN) architecture that has been proven to have better classification performance than other GNN architectures.

One of the first GNN architectures that was invented was basically dealing with the graph in a similar way as a CNN processes images or any kind of typical grid-structured data [26]. In the same way that CNN filters are convoluted with a portion of the input, the same applied to the Graph Convolutional Networks (GCN). The main difference lies in the fact that in GCNs the portion of the input is a subgraph (k-hop neighborhood) “around” a central node, whereas in a CNN the neighborhood of the central element is structured as a grid.

In figure 2, the CNN’s and GNN’s basic aggregation operation involving information from the neighborhood is depicted. Mathematically, this can be described by the following operation:

**Fig. 1.**
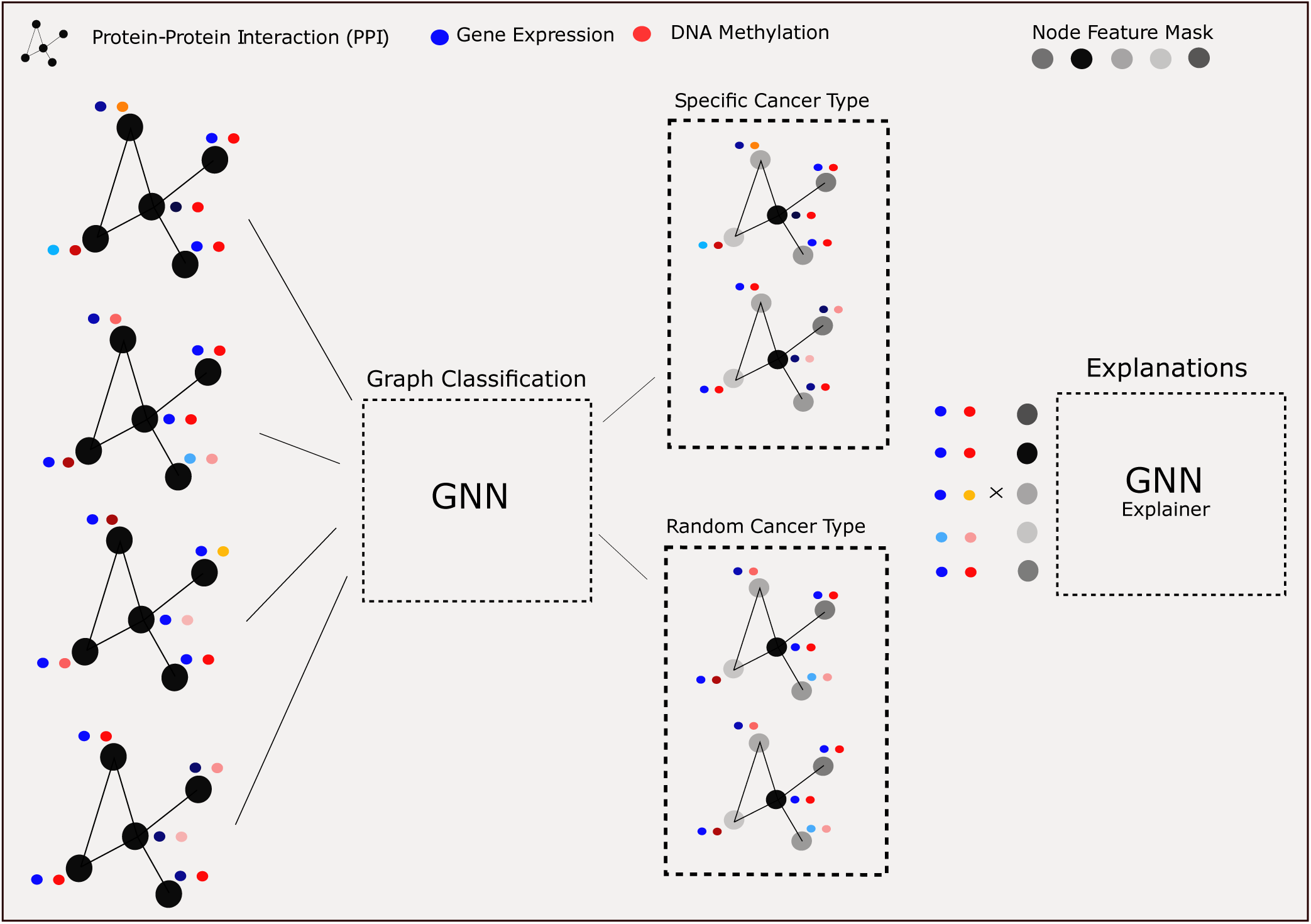
Illustration of patient classification into a cancer-specific and randomized cancer group using explainable Graph Neural Networks. Each patient is represented by the topology of an protein-protein interaction network (PPI). Nodes are enriched by multi-omic features from gene expression and DNA Methylation (colored circles). The topology of each graph is the same for all patients, but the node feature values vary, reflecting the cancer-specific molecular patterns of each patient.

**Fig. 2.**
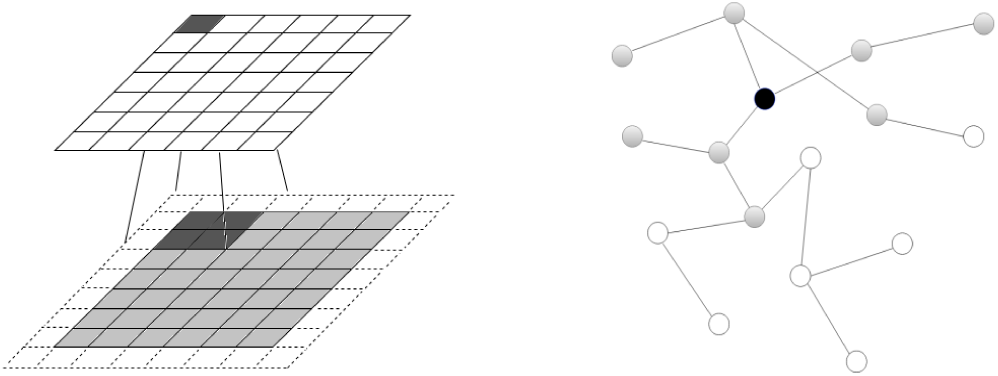
On the left side of the figure, the input and output grid of a two-dimensional convolution operation with a kernel of size 3 × 3, zero padding, and stride 1 is depicted. The kernel itself is not shown in the figures. The kernel slides over parts of the input grid. On the right side, the two-hop neighborhood of the centre node (marked with black color) is painted gray. The features of the neighboring nodes will be used to define updated values for the features of the currently processed node.

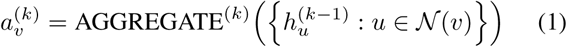

where *k* is the number of aggregation iterations and equals to the number of hops of the neighborhood 𝒩 that will be considered. The aggregation operation uses all features of the neighboring nodes (denoted with *u*); those can be of any form and can numerically encode several characteristics like size, color, shape and so on. As CNNs that process images typically aggregate three values (RGB) of each neighboring pixel, the GNN will aggregate any number of features representing any feature selected by the domain expert and the data scientist. Those features are denoted with *h* and are also called embeddings.

After the aggregation operation, the combine operation provides the new values for the features of node *v* 2:

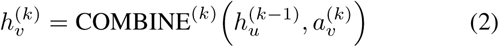

Those two operations, namely aggregate and combine are performed several times. The initial values of the features are replaced with new, informed ones that help the task at hand. Typically, GNNs can be used for node classification, link prediction and graph classification. Node as well as graph classification use the end values of the node features after the last application of aggregate and combine.

Until now, the way aggregate and combine are implemented is not fully addressed. The underlying operation in the GCN [26] is an element-wise mean pooling followed by a Rectified Linear Unit (ReLU) nonlinear activation function. The researchers that invented GIN [25] have proven that aggregations that are implemented by the `mean()` and `max()` function cannot distinguish between very simple graph structures; therefore, they are not adequate for computing embeddings, especially when the task is graph classification. Figure 3 shows three pairs of graphs that GNN architectures that use `mean()` and `max()` cannot differentiate. In the first pair, all nodes have the same values *h*_1_ in their features. In this case, both the mean and the maximum value over an arbitrary number of nodes in the neighborhood will be the same. In the second case, the maximum of *h*_1_, *h*_2_, *h*_3_ equals the maximum of *h*_1_, *h*_2_, *h*_3_, *h*_3_. By the same means, the maximum in the third case will fail for the same reason as in the second case. Furthermore, the mean will be the same because 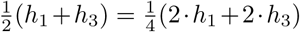.

**Fig. 3.**
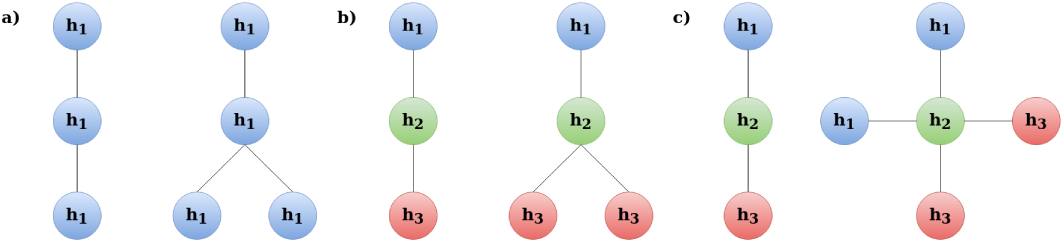
Three pairs of small graphs that GNN architectures that use `mean()` and `max()` functions for aggregation cannot differentiate.

Therefore, the researchers came up with the idea of using the `sum()` function as an aggregator. In all cases depicted in figure 3, the sum of the feature values of the graphs being compared is different. That shows the extended ability of the GIN architecture to discriminate more powerfully than any other GNN architecture.

As far as the combining step is concerned, the GIN architecture learns a function with the use of Multilayer Perceptrons (MLPs). This provides the necessary flexibility for injectiveness, maximum possible discrimination ability as well as the property of mapping similar graph structures to nearby embeddings. The overall equation of aggregation and combine steps in GIN is provided by the following equation 3:

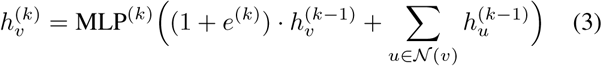

More information about the derivation of 3, as well as theoretical basis can be found in [25]. The exact architecture that was used for the proposed application consists of five Multilayer Perceptrons (MLPs). Each of those perceptrons is preceded with a pooling layer and succeeded with a batch normalization layer. Following those, there are five fully connected layers. Within the fully connected layers, each neuron is connected to all neurons in the layer before and after that layer. The MLPs consist of three layers, of which two are fully connected layers and layer for batch normalization layer. Batch normalization applies a transformation that maintains the mean output close to zero and the output standard deviation close to one. This technique helps to compensate the vanishing/exploding gradients problem to a certain extend [27].

### B. Explainable AI for disease subnetwork detection

In recent years, parallel to the development of different Graph Neural Network architectures, several strategies were invented to explain their decision process. Most of them are built on the assumption that a part of the input was the most important for the prediction in a similar way that the explanation for an image classification CNN will point out the areas in the image that were decisive and will ignore the ones that contain background. Explainable AI methods usually search for relevant subgraphs and their motifs [28], [29], walks [30] or even try to create Probabilistic Graphical Models [31], [32] which are causal structures, out of counterfactual examples that are computed by informed optimization problems [33].

GNNExplainer is used to compute the important subgraph *G*_*S*_ of the computation graph *G*_*c*_ of an input graph *G* that is going to be explained. This is achieved by graph masking as well as node feature masking, where the goal is to learn to mask the relevant part of the computation graph as well as the decisive node features. Those masks are found by an optimization algorithm that iteratively tries to find the substructure that maximizes the mutual information w.r.t. the prediction score. Equation 4 shows the optimization rule, where *X*_*S*_ is a subset of the the features of the nodes in the subgraph *G*_*S*_. **Y** represents the predicted label distribution; thereby the optimization process as a whole uses the change of the predicted label’s distribution as “guidance”

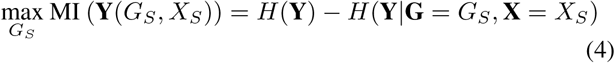

The random variables are denoted with bold letters, whereas instantiations (possible outcomes) thereof with non-bold letters.

In this work we employ a simplified version of the GNNexplainer with an induced sampling scheme. Since we apply the explanations on a graph classification task, where all graphs have the same topology, the equation 4 can be re-written as

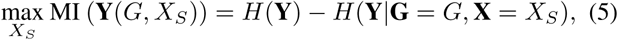

where G is the original graph. We employ a node mask on **X** such that a subset of nodes in *X*_*S*_ can be inferred to maximize mutual information with a minimal set of features. To make the optimization process more efficient and tractable, the researchers came up with several constraints and improvements. For more details please see [28]. Accordingly, we solve the following optimizing function by gradient decent.

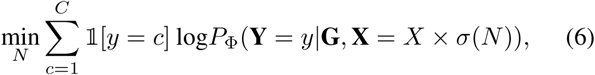

where *N* ∈ ℝ^1×#*V*^ specifies the learned feature node mask passed through the sigmoid function *σ. X* × *σ*(*N*) is the row-wise multiplication of *X*, where the rows reflect the nodes and the columns are representing the features. The number of nodes in *X* can be constrained; this is a configurable parameter in the implementation provided in the github repository. In the above expression c denotes one of the possible C classes of a classification class.

GNNexplainer allows for node feature masks as well es edge masks. The GNNexplainer, however, may not be applicable for *model-wide* explanations [29]. This is due to the fact that it optimizes a specified node and edge mask with regard to a single input graph. As a consequence, explanations may not reflect the *global* decisions made by the GNN classifier [29]. In fact, the mentioned problematic was recently addressed by a method called PGExplainer [29]. However, the PGExplainer explicitly works on edge masks and thus requires the GNN model to internally adjust edge weights, which was not applicable in our case. Thus, we propose a slight modification of the GNNexplainer for *model-wide* explanations.

We randomly sample graphs from the input space, while optimizing one single node feature mask *N* ∈ ℝ^1×#*V*^. After a certain number of epochs the sampling scheme is repeated. As a result, the values of the node feature mask converge to *global* node importance values. This approach is very much related to classical feature selection. Instead of inferring explanations for a single instance, we provide feature importance values for the whole set of samples. The proposed important node attributes may be an efficient technique for GNN-based dimension reduction, so that reduced subnetworks could provide more parsimonious models which to this end may generalize better on unseen test data.

Ultimately, disease subnetworks are detected with the following approach. First, we assigned edge relevant scores by calculating the average node feature importances of two connected nodes, inferred by our modified GNNexplainer. The obtained edge-specific scores are used to weight the edges of the PPI network. Following, a louvain method [34] for community detection was applied to the weighted input graphs. The detected communities are ranked according to their average edge importance scores. The top-ranked community represents the detected disease subnetwork.

### C. Sanity checks on synthetic data

We have validated our approach on synthetic Barabasi networks. We have generated 1000 networks comprising of 30 nodes each. Node feature values were generated from a normal distribution with *N* (*μ* = 0, *σ*). For each of the 500 networks we sampled two node feature values from *N* (*μ* = −1, *σ*), and *N* (*μ* = 1, *σ*) respectively for the other 500 networks. We assigned these feature values to two randomly selected connected nodes. We have evaluated whether and to what extend the GNNexplainer explanations successfully uncover the selected edge and the corresponding nodes. We varied the *σ* values and on the stability and robustness of the explanations. Results of these sanity checks are shown in Figure 5 and Table I of the ‘Materials and Methods’ Section.

**TABLE I.** Performance of the GNN classifier and its explanations

| Noise ($\sigma$) | Accuracy of GNN classifier | Coverage of GNNexplainer |
| --- | --- | --- |
| $\sigma = 0.1$ | 100% | 1 |
| $\sigma = 0.3$ | 98% | 1 |
| $\sigma = 0.5$ | 91% | 0.94 |
| $\sigma = 0.7$ | 79% | 0.75 |
| $\sigma = 1$ | 70% | 0.44 |

### D. TCGA human cancer data

We have downloaded molecular multi-modal data from the *linkedomics*.*org* server [35]. The authors provide harmonized multi-omics data sets retrieved from The Cancer Genome Atlas (TCGA) database (https://www.cancer.gov/tcga), which represents one of the largest collections of multi-omics data sets. It contains molecular and genetic profiles for over 33 different cancer types from 20,000 individual tumor samples [36]. Here, we analyzed three different cancer types, namely Kidney Renal Clear Cell Carcinoma (KIRC), Breast Invasive Carcinoma (BRCA), and Lung Adenocarcinoma (LUAD), for which we detect cancer-specific subnetworks which are substantially different from a randomized control group. The control group consists of 200 randomly sampled patients across randomly selected cancer types.

We have downloaded gene expression (HiSeq) and DNA methylation (HM450K) to enrich the nodes with feature values. We harmonized the data sets such that for every patient multi-source informations is available. Furthermore, we filtered for cancer-relevant genes as proposed by [8]. Genes with missing values at least for one patient were removed from the analyses. The obtained numerical data matrices were normalized using min-max normalization.

The protein-protein interaction network was retrieved from the STRING database [37]. We only kept nodes for which both, mRNA gene expression data and DNA methylation data was available. We deleted edges with relevance scores lower than the 95-percentile. In case this filtering resulted in a multi graph, we kept the sub-network with the highest number of nodes.

## IV. Results

### A. Synthetic data sets

Results obtained from synthetic data indicate that the proposed GNNexplainer node feature mask successfully uncovers the GNN black-box decisions. Figure 4 shows an example of a simulated Barabasi graph whose node features are generated with *N* (*μ* = 0, *σ* = 0.1). The graph consists of 30 nodes and 209 edges. We simulated 1000 graphs with that exact same topology with varying the node feature values. The features values of the selected edge 4-5 was generated from *N* (*μ* = − 1, *σ* = 0.1) for 500 graphs, and *N* (*μ* = 1, *σ* = 0.1) for the other 500 graphs. The sampling induced variation of the GNNexplainer detects the selected edge, inferring the highest score of 0.95 for it (see Figure 4. Notably, the selected edge is detected even though it is not placed within a highly connected community. This observation suggests that the GNN classifier as well as the explainer are not biased towards nodes with high edge degree.

**Fig. 4.**
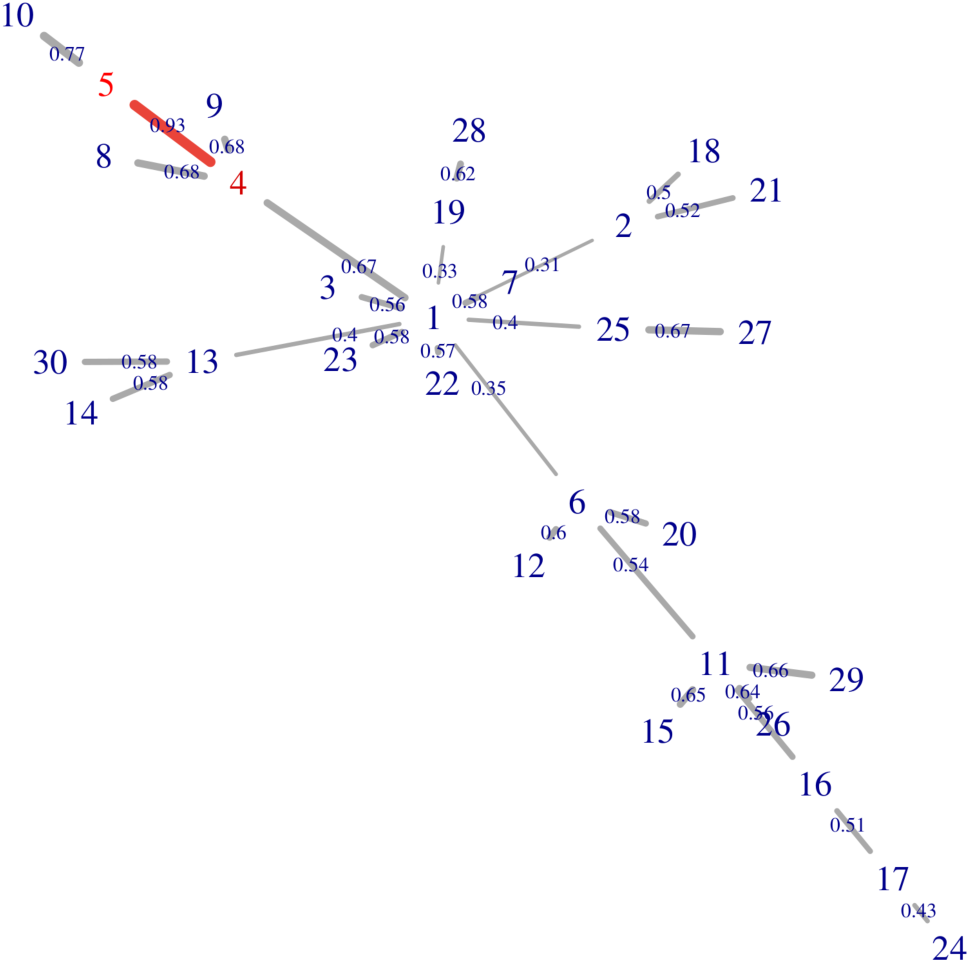
Barabasi graph. Shown is a simulated barabasi graph with 30 nodes and 29 edges. The edges are labeled by the importance scores obtained from our modified GNNexplainer. Selected edge is 4-5 with the highest score of 0.93.

We repeated the analyses with varying graph topologies and variable *σ* values of the selected edge node features. As can be seen from Figure 5, the selected edge is within the top-2 ranked edges in all cases, when *σ* values are lower than 0.5. For *σ >* 0.5, the accuracy of the GNN classifier reduces significantly, and explanations get worse accordingly. For *σ* = 0.3 a single outlier run with low coverage can be observed. However, the median coverage values are still at 100%. Table I shows the median coverage values for the top-1 ranked edges as well as the accuracy of the GNN classifier. As expected, the more noise it added to the synthetic data, the lower is the accuracy of the GNN classifier.

**Fig. 5.**
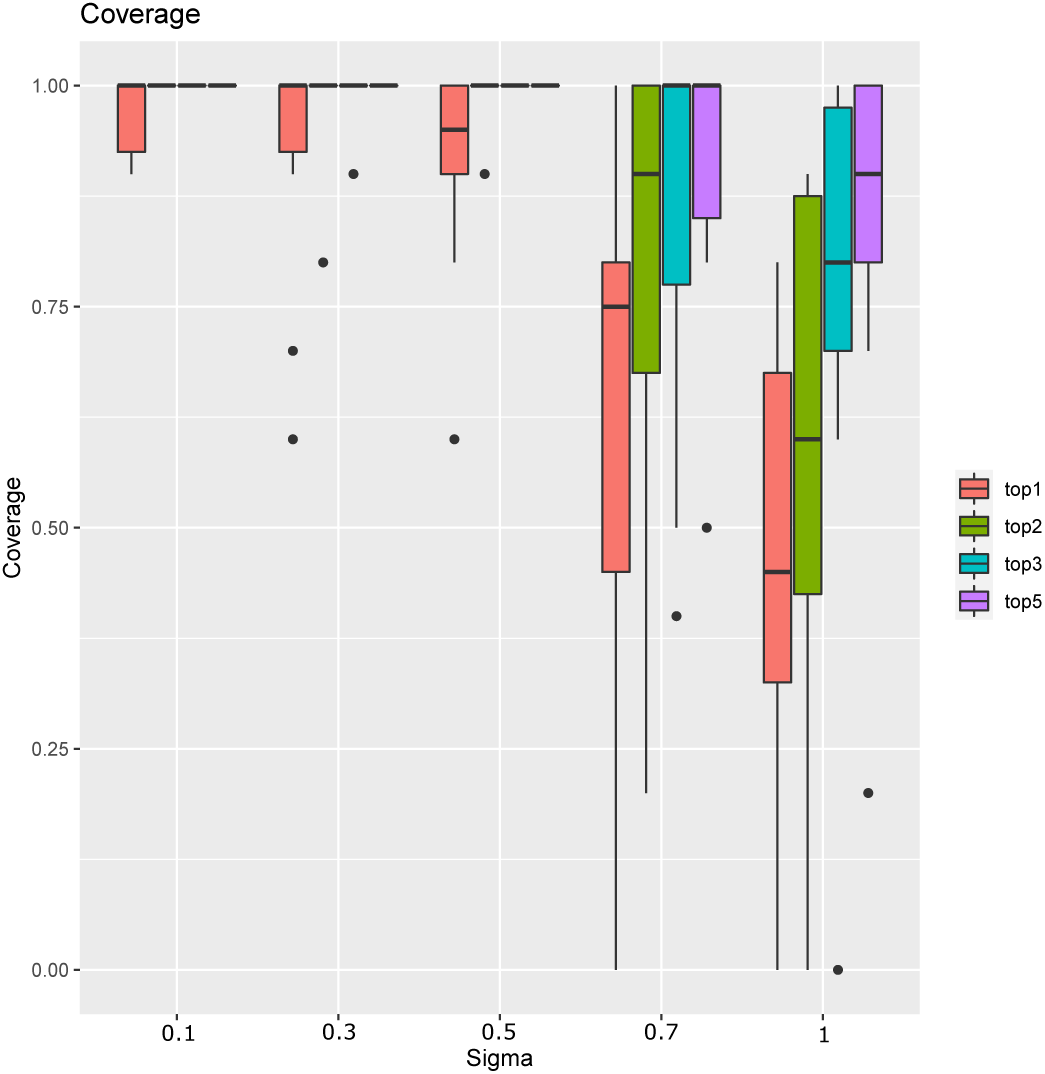
Results on synthetic Barabasi graphs. Shown is the coverage, which is measured by the number of times the selected edge is ranked within the top-k elements. The edge importance values are calculated based on the GNNexplainer with induced sub-sampling.

Interestingly, explanations in the case of *σ* = 0.3 are at 100% while the overall accuracy of the GNN model is at 98%. The better accuracy of the explanations is due to the fact that the GNNexplainer perturbates the features such as the predicted class becomes more likely. We believe that our modified GNNexplainer can also be used as a dimension reduction algorithm, where the most important nodes are filtered and a new classifier is trained on this reduced set. Further investigations are needed to elaborate on this potential capacity.

### B. Application to TCGA cancer data

The accuracy of the employed GNN model on the cancer data sets is shown in Table II. The models were trained using a validation set, and we applied an early stopping criteria. Test performance is reported based on a 80% − 20% train-test data split. The number of epochs was set to 20, and we kept the model with lowest *loss* value on the validation set. This model was finally applied to the hold-out test data set.

**TABLE II.** Accuracy of the GNN classifier on TCGA cancer data

| Cancer type | mRNA | DNA Methylation | Multi-Omics |
| --- | --- | --- | --- |
| KIRC | 0.63 | 0.75 | 0.89 |
| BRCA | 0.54 | 0.66 | 0.77 |
| LUAD | 0.60 | 0.91 | 0.88 |

We could observe that in two out of three cases incorporating multiple biological sources was beneficial. An exception was LUAD. The GNN model performed best when it was trained exclusively on DNA Methylation data (see Table II). Interestingly, mRNA as a single-source node feature performed worse than DNA Methylation for all analyzed cancer types. This observation might indicate that KIRC, BRCA, and LUAD have similar mRNA gene expression levels, but differ mostly due to epigenetic factors. It would require a further in-depth analysis in order to proof this hypothesis, and this is out of scope for this article. Overall, the performance was best for the KIRC data set, with an accuracy of 0.89, for which we did further investigations.

We applied our modified GNNexplainer to the KIRC GNN model in order to verify the most important network regions for classification. From initially 2049 genes and 13588 edges, we have detected 36 modules in total. The top-ranked module, according to its average edge importance score, is shown in Figure 6. The module consists of eight genes, namely GLE1, POLR2A, SMO, SORL1, TNS1, TRIM25, UBR5, and UTRN. These genes are connected by 22 edges. The genes SORL1, SMO, and POLR2A have the highest connectivity with four edges, where POLR2A has the highest average edge importance with 0.88. The POLR2A gene is essential for cell survival, and is almost always co-deleted with TP53, in many human cancers, including colorectal, breast, ovarian, kidney, liver, and pancreatic cancer [38]. POLR2A was very recently indentfied as a potential binding protein POLR2A of ARGLU1, a potential therapeutic target gene [39].

**Fig. 6.**
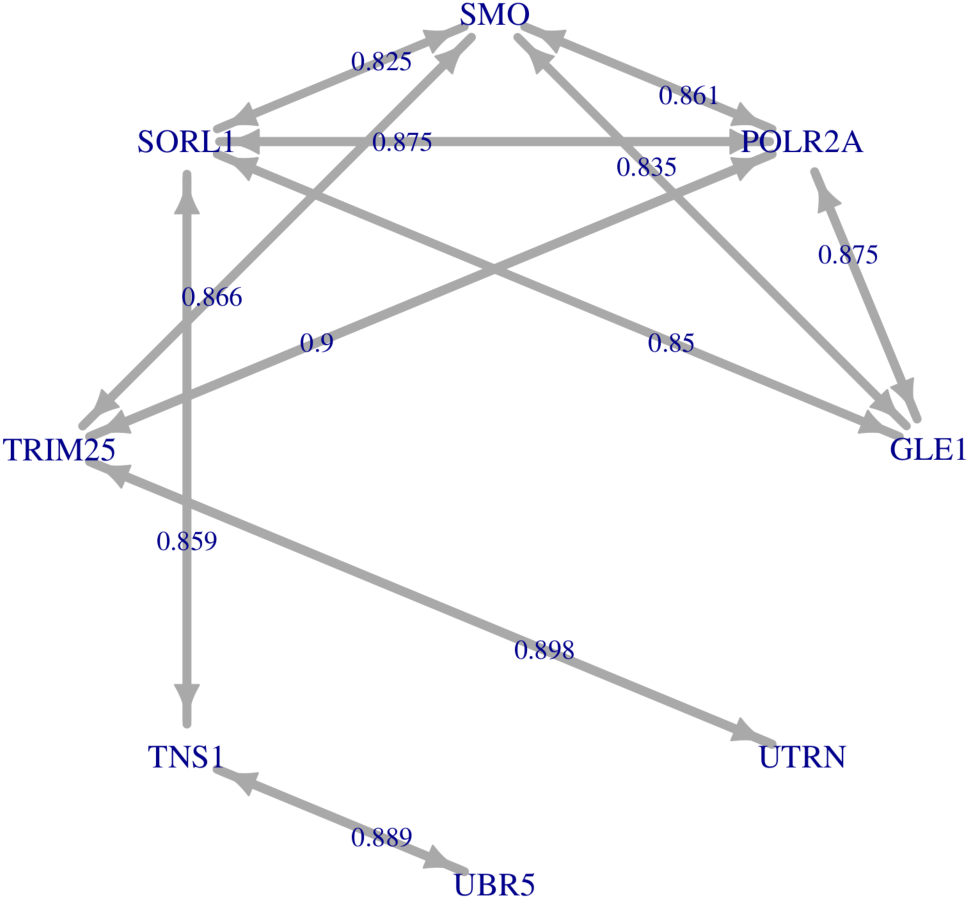
Kidney cancer disease subnetwork. Shown is the top-ranked community inferred by our proposed explainable GNN pipeline. Edge weights are calculated using the GNNexplainer with induced sub-sampling.

As can be seen from Figure 6. The edge between TRIM25 and POLRS2A has the highest edge importance with 0.9. TRIM25 has an important RNA-binding role in antiviral defense [40]. We did not find any indication of a special importance of the interactions between POLR2A and TRIM25 within the current cancer literature. However, the fact that both are binding genes makes an in-depth analyses intriguing.

## V. Discussion and Conclusion

In this article we have introduced a framework for the detection of disease subnetworks. We argued that the incorporated PPI knowledge-graph restricts the deep model to learn on more reliable and biological meaningful trajectories compared to classical deep learning approaches. This is important and it may fasten the way towards the discovery of novel biomarker.

We have introduced a simple modification of the GNNexplainer program such that it computes model-wide explanations. This was realized by randomly sampling networks from the input space, while optimizing a single node mask. From this node mask edge relevance scores were computed and were assigned as edge weights to the PPI network. Disease subnetworks are finally inferred by a weighted community detection algorithm.

We have demonstrated PPI disease subnetwork detection from patients suffering kidney cancer. Each patient was modeled as a PPI network comprising multi-omic node features from mRNA gene expression and DNA methylation data.

Finally, we have implemented our proposed methodology within the GNN-SubNet program. We plan to develop this program further and add features from various angles. For instance, several additional GNN-based explainer will be incorporated. Explainer like PGM-Explainer and GNN-LRP are of particular interest. The PGM-Explainer is able to demonstrate the dependencies of explained features in form of conditional probabilities [33], which may contribute to better causal understanding of the explanations. The GNN-LRP method explains the GNN classifier using higher-order expansions [30]. A main advantage compared to other methods is that it is capable of reporting on both, positive contribution as well as negative contribution of features to a particular prediction. All this together could help to increase the interpretability of the detected disease subnetworks.

The explanations of different explainer may differ with regard to their quality and the introduced application. An indepth comparison is needed which to this end may help for more specialized methods tailored towards a specific application domain.

There are many opportunities for future work. Biomedical experts are increasingly faced with high-dimensional data sets, accelerated by the trend toward precision medicine. Although human experts excel at pattern recognition in dimensions of less than three, most biomedical data exist in dimensions much greater than three, making human manual analysis difficult, even virtually impossible. Biomedical experts are therefore less and less able to deal with such data in their daily routine, which requires efficient, usable, and useful methods, algorithms, and tools to interactively gain insight into such data. A synergistic combination of graph theory [41], topological [42] and entropy [43] analysis methods seems to be very promising for the future [44].

## Abbrevations

CNN: Convolutional Neural Network
GNN: Graph Neural Network
GIN: Graph Isomorphism Network
MLP: Multi Layer Perceptron
AI: Artificial Intelligence
XAI: Explainable Artificial Intelligence
DT: Decision Tree
DF: Decision Forest
TCGA: The Cancer Genome Atlas
PPI: Protein-Protein Interaction Network
KIRC: Kidney Renal Clear Cell Carcinoma
BRCA: Breast Invasive Carcinoma
LUAD: Lung Adenocarcinoma
ReLU: Rectifiec Linear Unit

## VI. Author Contributions

BP designed the project, developed the methods, and performed the analyses. BP and AS implemented the proposed framework in python. BP, AS, and AH wrote the manuscript. AH raised the funding. All authors reviewed the article.

## VII. Data and Code Availability

The results shown here are in whole or part based upon data generated by the TCGA Research Network: https://www.cancer.gov/tcga. Python Code is freely available on GitHub (https://github.com/pievos101/GNN-SubNet), including example files and documentation.

